# Transposition of *HOPPLA* in siRNA-deficient plants suggests a limited effect of the environment on retrotransposon mobility in *Brachypodium distachyon*

**DOI:** 10.1101/2023.09.25.559196

**Authors:** Michael Thieme, Nikolaos Minadakis, Christophe Himber, Bettina Keller, Wenbo Xu, Kinga Rutowicz, Calvin Matteoli, Marcel Böhrer, Bart Rymen, Debbie Laudencia-Chingcuanco, John Vogel, Richard Sibout, Christoph Stritt, Todd Blevins, Anne C. Roulin

## Abstract

Long terminal repeat retrotransposons (LTR-RTs) are powerful mutagens regarded as a major source of genetic novelty and important drivers of evolution. Yet, the uncontrolled and potentially selfish proliferation of LTR-RTs can lead to deleterious mutations and genome instability, with large fitness costs for their host. While population genomics data suggest that an ongoing LTR-RT mobility is common in many species, the understanding of their dual role in evolution is limited. Here, we harness the genetic diversity of 320 sequenced natural accessions of the Mediterranean grass *Brachypodium distachyon* to characterize how genetic and environmental factors influence plant LTR-RT dynamics in the wild. When combining a coverage-based approach to estimate global LTR-RT copy number variations with mobilome-sequencing of nine accessions exposed to eight different stresses, we find little evidence for a major role of environmental factors in LTR-RT accumulations in *B. distachyon* natural accessions. Instead, we show that loss of RNA polymerase IV (Pol IV), which mediates RNA-directed DNA methylation in plants, results in high transcriptional and transpositional activities of RLC_BdisC024 (*HOPPLA*) LTR-RT family elements, and that these effects are not stress-specific. This work supports findings indicating an ongoing mobility in *B. distachyon* and reveals that host RNA-directed DNA methylation rather than environmental factors controls their mobility in this wild grass model.

**Author summary:** Long terminal repeat retrotransposons (LTR-RTs) are major components of plant genomes. Their ‘copy-and-paste’ replication mechanism allows them to rapidly increase in copy number, with potentially negative effects on host fitness. On the other hand, because they can rewire transcriptional networks and alter phenotypes, their mobility is an important driver of evolution. Ever since their discovery, LTR-RT activity has been linked to stress exposure, suggesting that LTR-RTs modulate the pace of evolution in response to the environment. In this study, we test this hypothesis by harnessing the genetic variation in a set of 320 natural accessions of the Mediterranean grass *Brachypodium distachyon* originating from diverse habitats. We find little evidence for the importance of stresses in activating *B. distachyon* LTR-RTs. Instead, we show that the loss of RNA polymerase IV, a component of plant retrotransposon silencing, leads to the activation and transposition of an LTR-RT family that we name *HOPPLA*. *HOPPLA* is the first LTR-RT family in *B. distachyon* shown to transpose in real-time. These findings open up new avenues for studying retrotransposon-mediated evolution in this close relative of staple crops, such as rice and wheat.

## Introduction

Transposable elements (TEs) are DNA sequences with the ability to form extrachromosomal copies and to reintegrate elsewhere into the host genome. In plants, TE-derived sequences are ubiquitous and can constitute more than 80 % of the genome [1]. In addition to playing a major role in genome size variation (e.g. [2–5]), TEs can alter gene expression by acting as promoters or by providing *cis*-regulatory elements to flanking regions [6–8]. TEs are therefore a major source of genetic change. Given that they are more likely than classic point mutations to cause extreme changes in gene expression and phenotypes [9–11], they might be especially useful when the survival of an organism or its descendants depends on a quick response to new or challenging environmental conditions (for review [12–15]). Paradoxically, while population genomics data have revealed ongoing TE activity in natural plant populations (e.g. [16–18]), only a handful of TE families have been experimentally shown to transpose. Therefore, how often or under which natural conditions TEs are activated in the wild remain open questions. In addition, while ongoing transposition is essential for TEs to survive, the presence of mobile and potentially ‘selfish’ DNA sequences requires the host to evolve robust silencing mechanisms to prevent an uncontrolled TE proliferation. TE activity thus remains a major puzzle in the field of evolutionary genomics.

In plants, the defence against TEs is multi-layered, comprising repressive histone modifications, DNA methylation and RNA interference [19–21]. One of the main players of TE silencing is the RNA-directed DNA methylation (RdDM) pathway, which involves two plant specific RNA-polymerases derived from Pol II, namely Pol IV and Pol V. The largest subunits of each polymerase (NRPB1, NRPD1 and NRPE1, respectively) assemble with other proteins into enzymes with distinct RNA products and functions [22,23]. As a core component of RdDM, Pol IV (including NRPD1) transcribes TE regions into the precursors of functionally specialized 24 nt small interfering RNAs (siRNAs) [21,23,24]. Upon the base pairing of 24 nt siRNAs to scaffold transcripts produced by Pol V, the *de novo* DNA methyltransferase DRM2 [25] is recruited to mediate the methylation and subsequent transcriptional repression of TEs. The essential role of RdDM in TE silencing has been shown in *A. thaliana*, where the knockout of Pol IV and resulting depletion of 24 nt siRNAs leads to a drastically increased heat-dependent transposition of the *ONSEN* family [26,27].

The case of the heat-responsive *ONSEN* elements not only illustrates the importance of epigenetic silencing in regulating TEs but also demonstrates that environmental factors may modulate the dynamics of TEs in plants. Since their discovery by Barbara McClintock, who linked the mobility of Ac/Ds elements in maize to the occurrence of a ‘genomic shock’ [28], the activity of TEs has been frequently associated with the presence of biotic and abiotic stressors. In fact, certain TEs can sense specific physiological states of their host and use them to initiate their own life cycle [29]. Besides *ONSEN* in *A. thaliana* [30], the cold inducible *Tcs1* element in blood oranges [8], *Tos17* that gets activated during tissue culture in rice [31], *Tnt1* of tobacco that reacts to wounding [32] or the drought responsive *Rider* retrotransposon in tomato [33] are other prominent cases of stress-responsive TEs in plants. Mechanistically, stress can activate TEs via specific motifs allowing the binding of transcription factors and the subsequent recruitment of the transcription machinery to their promoter-like sequences [18,30,34,35]. The small window of increased activity during well-defined physiological states suggests that some TEs have evolved a distinct lifestyle or ‘niche’ to successfully reproduce [36–39].

Since transcription constitutes the initial step to transposition, the abundance of TE transcripts is often used as a proxy for TE activity [40]. However, the life cycle of TEs is complex [41] and the fate of TE transcripts depends on many factors. For instance, several transcriptionally active TEs have accumulated mutations that prevent the production of enzymes needed for their autonomous transposition [42]. To selectively capture TEs that are not only transcriptionally active but also capable of transposing, several protocols have been developed, including ALE-seq [43], VLP-seq [44] and mobilome-seq [45]. These recent approaches have been particularly successful when targeting long terminal repeat retrotransposons (LTR-RTs), which represent the largest fraction of TE-derived sequences in plant genomes [46]. Indeed, LTR-RTs transpose through a copy-and-paste mechanism which involves the reverse transcription of a full-length RNA intermediate [41,47]. As part of their life cycle and presumably through auto-integration, non-homologous and alternative end-joining, active LTR-RTs can also form extrachromosomal circular DNA (eccDNA) intermediates [44,47–51], whose detection by mobilome-seq can be used as a proxy for their mobility [45]. For instance, mobilome-seq has been successfully used to track full-length eccDNA of mobilized autonomous RTs, containing both LTRs (2-LTR circles), in plants such as *A. thaliana* and rice [45,52].

While the activity of LTR-RTs has been extensively studied in the model *A. thaliana* [18], the interplay between genetic and environmental factors in other wild plant species remains poorly investigated. To clarify these questions, we exploit here the *Brachypodium distachyon* diversity panel [53] and explore LTR-RT activity in a wild monocot*. B. distachyon* is a Mediterranean grass with a compact diploid genome of ∼272 Mb [54,55] harboring 40 LTR-RT families [56] that constitute about 30 % of the genome [55]. In *B. distachyon*, LTR-RTs not only evolved varying insertion site preferences [56] but also significantly differ in terms of transposition dynamics [17,56]. While we previously suggested an ongoing transposition of LTR-RTs based on population genomics data [17,57], here we aimed to clarify the contribution of genetic and environmental factors to LTR-RT activity in *B. distachyon*. To that end, we combined population genomics data available for 320 natural accessions with mobilome-seq under different stress conditions and asked: (i) does the accumulation of LTR-RTs in these natural accessions correlate with environmental variables, (ii) is LTR-RT mobility induced by specific stresses, and (iii) which genetic factors influence the accumulation of LTR-RTs?

## Results

### Abundances of LTR-RT families differ but show limited association with bioclimatic variables

*B. distachyon* naturally occurs around the Mediterranean rim (**Fig 1A**) and groups into three main genetic lineages (A, B and C) that further split into five genetic clades [53,58]: an early diverged C clade and four clades found in Spain and France (B_West), Italy (A_Italia), the Balkans and coastal Turkey (A_East) and inland Turkey and Lesser Caucasus (B_East). We first aligned genomic reads of 320 accessions to the LTR-RT consensus sequences of *B. distachyon* obtained from the TRansposable Elements Platform (TREP). We then computed the abundance of TE-derived sequences, hence a proxy for copy numbers (pCNs), for the 40 annotated LTR-RT families using a coverage-based approach accounting for sample sequencing depth (see Materials and Methods). We favored this approach over an analysis based on transposon insertion polymorphisms (TIPs) because estimates based on TIPs are reference genome-dependent and biased by the phylogeny in our study system. We have, for instance, previously shown that accessions from the B_East clade harbor significantly less TIPs than accessions from the A_East clade due to the fact that the reference genome Bd21 belongs to the B_East clade [17]. In addition, whole-genome *de novo* assembly of 54 *B. distachyon* natural accessions and the subsequent pangenome analysis revealed that non-reference accessions display large genomic variations [59], which may further bias the estimates of TIP abundances.

**Fig 1.**
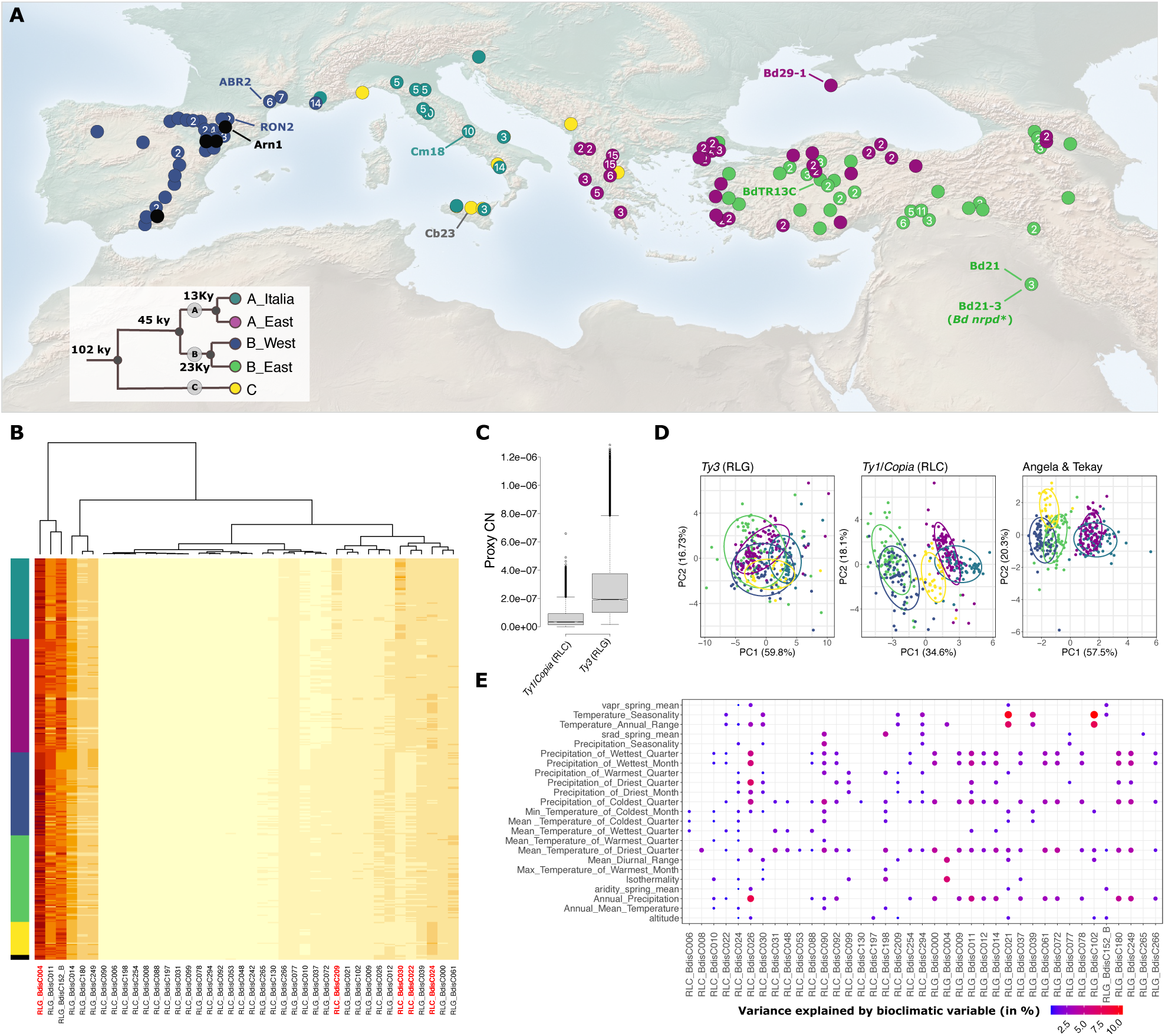
Natural diversity of proximal copy number (pCN) variation of LTR-RTs in *B. distachyon*. (**A**) Origin of the 320 natural accessions included in this study. Accessions that were used for the mobilome-seq are labelled in the map and a PCA with the bioclimatic variables of their place of origin is depicted. Colors of points correspond to the genetic clades whose estimated split is shown in the phylogenetic tree. Black points indicate that the accession cannot be clearly assigned to one genetic clade. Numbers in dots indicate how many sequenced natural accessions were sampled in the marked area. The map has been obtained from (https://www.naturalearthdata.com/downloads/50m-cross-blend-hypso/50m-cross-blended-hypso-with-shaded-relief-and-water/). (**B**) Heatmap with read counts of TE-derived sequences (proxy for the copy number (pCN) variation) of all 40 annotated LTR-RTs in 320 natural accessions of *B. distachyon.* TE-families are clustered by their pCNs (dendrogram above the heatmap) and accessions are sorted according to their phylogeny. Names of recently active TE-families are highlighted in red. (**C**) Overall estimates of the copy numbers of *Ty1/Copia* and *Ty3*-type LTR-RTs in 320 natural accessions. (**D**) PCAs of pCNs of all *Ty3*, *Ty1/Copia* and all recently active LTR-RT families, highlighted in (**B**) belonging to the Angela & Tekay families (RLG_Bdis004, RLC_BdisC030, RLC_BdisC209, RLC_BdisC024 and RLC_BdisC022). Colors of points indicate the genetic clade of accessions. (**E**) Output of the LMM analyses between pCNs of LTR-RTs and bioclimatic variables at the accessions’ origin. Bubbles indicate a significant association (P-value <0.05). Colors and sizes of bubbles show the part of the variance (marginal R^2^) explained by the bioclimatic variables in %.

The heatmap produced based on pCN variation (**Fig 1B**) showed that LTR-RTs underwent different transposon accumulations. We found that *Ty3* elements (RLG) harbor higher pCNs than *Ty1/Copia* (RLC) elements (Wilcoxon test, W = 6517781, p-value < 2.2e-16; **Fig 1B and 1C**). Furthermore, a PCA based on RLG pCNs did not allow us to discriminate accessions based on their phylogenetic relationship, while a PCA performed with RLC elements separated the samples by genetic lineage (**Fig 1D**). The strongest result was found with a PCA performed with the five youngest and putatively most recently active LTR-RT families found in the pangenome of *B. distachyon* (the Angela families RLC_BdisC022, RLC_BdisC024, RLC_BdisC030, RLC_BdisC209 and the Tekay family RLG_BdisC004) [56]), with the first and the second principal components together explaining more than 77.8% of the variance. Finally, with the exception of samples from the most recently diverged clades A_East and A_Italia (13 kya), they further allowed us to discriminate samples based on the genetic clade of origin (**Fig 1B and 1D**).

To test whether the accumulation of LTR-RT sequences correlated with environmental factors, we retrieved bioclimatic variables comprising precipitation, temperature, aridity levels, solar radiation and atmospheric pressure at each locality. We then ran linear mixed models (LMM) where pCNs per LTR-RT family was entered as the response variable, the bioclimatic variables entered separately as fixed factors and the clade of origin as random factors to account for population structure. Marginal R^2^ extracted for each LMM did not exceed 10% even for the putatively most recently active LTR-RT families (**Fig 1E**), indicating that albeit significant, the association between pCNs per family and the environment was mild in our study system. We observed similar associations between pCNs and bioclimatic variables when not accounting for population structure and running classical linear model analyses (**S1 Fig**). With the exception of the two relatively young and low-copy families RLC_BdisC010 and RLG_BdisC265 [56], for which more than 40 % of the variance in pCNs was explained by environmental factors, the LTR-RT families showed non-significant to mild associations with environmental variables (**S1 Fig**).

### Recently active LTR-RT families produce eccDNAs

The coverage-based approach for estimating LTR-RT pCNs considers all TE-derived sequences and does not take into account that individual families differ in their age structures, turnover times and proportion of full-length, potentially autonomous mobile elements [56]. We therefore complemented our *in silico* analysis by experimentally testing whether LTR-RT families were indeed still active *in planta* and whether the mild but significant correlation we observed between global variations in pCNs and bioclimatic variables may be due to a stress-specific activity.

To cover a wide range of the genetic, geographic and bioclimatic diversity of *B. distachyon*, we selected nine natural accessions belonging to the five genetic clades and originating from contrasting habitats (**Fig 1A**). Considering the role of Pol IV in the silencing of TEs in plants, we also included two independent Pol IV mutant lines, a sodium azide mutagenized line (hereafter *Bd nrpd1-1*) and a T-DNA line (hereafter *Bd nrpd1-2*), both carrying a homozygous mutation in the largest subunit of Pol IV (NRPD1; Bradi2g34876) in the Bd21-3 background. We exposed plants to eight different stresses, namely cold, drought, heat, salt, submergence, infection with *Magnaporthe oryzae*, treatment with glyphosate and chemical de-methylation and performed mobilome-seq on all resulting 105 samples (S6 **Table)**. Mobilome-seq was developed to specifically capture circular extrachromosomal DNA that is formed as an intermediate during retrotransposon mobility [45] (see Material and Methods).

The rolling circle amplification of eccDNAs during mobilome-seq results in a high coverage of mobile elements allowing to reconstruct near complete mobile LTR-RTs from short reads [45]. Following the removal of organelle-derived reads, we thus first assembled mobilome reads and only aligned the resulting ten longest contigs of each sample to the reference assembly of *B. distachyon*. The size selection of contigs allowed us to only retain the most relevant eccDNA candidates of each sample, potentially representing full-length mobile LTR-RTs. We then screened genomic regions for which at least three assembled mobilome contigs of different samples aligned and further assessed in which genotypes or stresses those contigs occurred. In a next step, we attempted to identify those genomic regions that were enriched in contigs assembled from mobilome reads from certain genotypes or stress conditions. We hence only retained circle-forming regions with a specificity above 50 % (i.e., regions for which more than half of the contigs belonged to a certain stress or genotype). In addition, we also kept recurrently active regions, present at a high frequency independently of the stress or the genotype in at least ten samples. In total, we retained 15 circle-forming regions, all of which contained TE sequences (**Fig 2**). Eight of these corresponded to the Angela family (RLC_BdisC024, RLC_BdisC022, RLC_BdisC030, RLC_BdisC209), four to CRM elements (RLG_BdisC039, RLG_BdisC102), and one each to the SIRE (RLC_BdisC026), the Alesia (RLC_BdisC010) and the non-autonomous and unclassified RLG_BdisC152 family. We hereafter refer to RLC_BdisC024 as *HOPPLA* (German allusion for the surprising finding of a jumping element). We did not find stress specificity in the formation of eccDNAs (**Fig 2A**). However, our results pointed to a genotype-dependent formation of eccDNAs for the RLC_BdisC209, RLC_BdisC026 and *HOPPLA* families (**Fig 2B**). In particular, two contigs containing *HOPPLA* elements were exclusively detected in the two *pol IV* mutants (**Fig 2B**).

**Fig 2.**
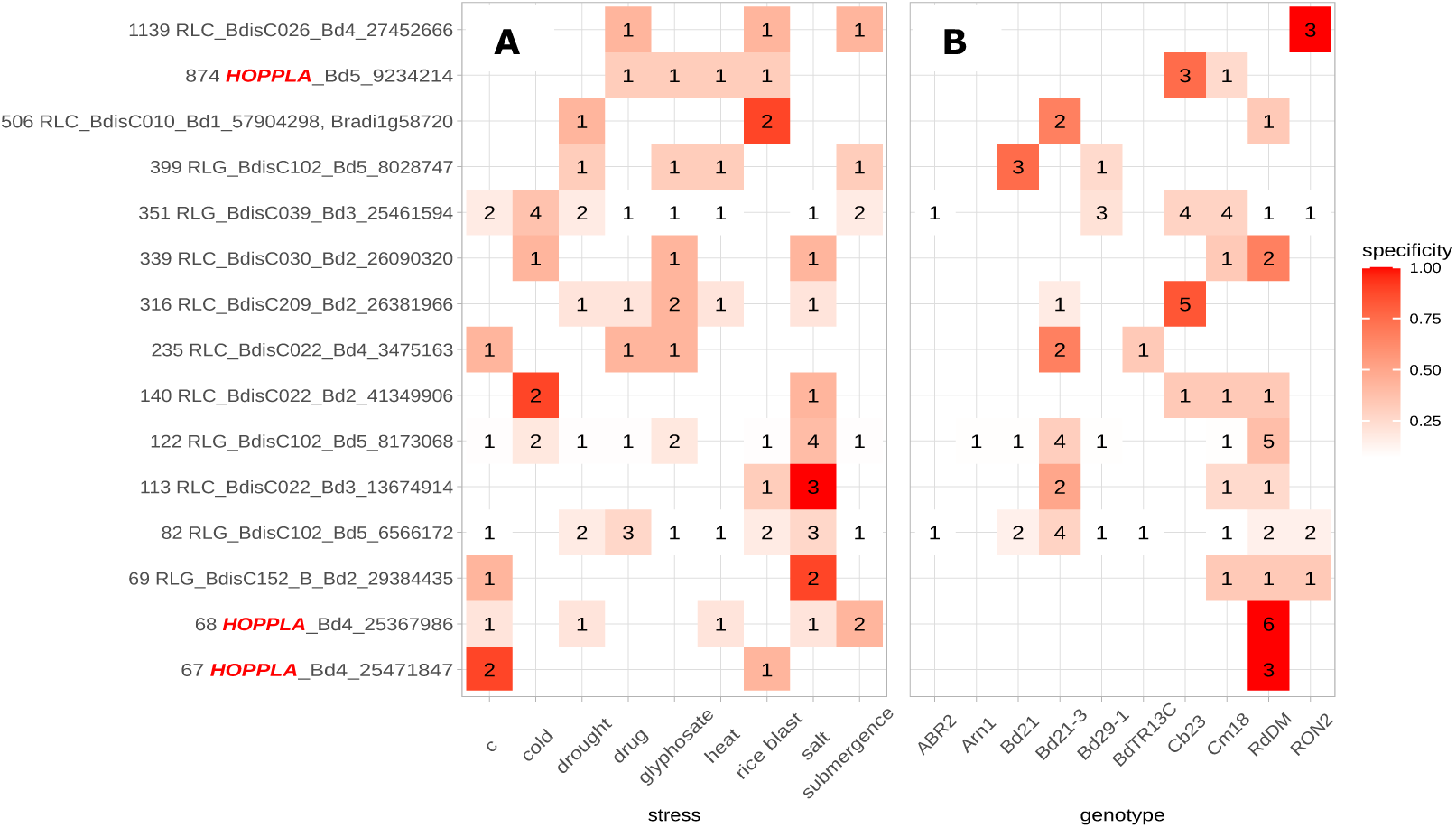
Assessment of LTR-RT mobility in *B. distachyon*. Stress (**A**) and genotype (**B**) specificity of the formation of eccDNA as determined by the alignment of assembled mobilome-seq reads. The color represents the degree of specificity and numbers indicate the count of samples from which one of their ten longest contigs aligns to each of the circle-forming regions. Annotations of regions are indicated on the y-axis. Loci containing *HOPPLA* (RLC_BdisC024) are highlighted in red. Multiple annotations in the same circle-forming region were concatenated. The two *pol IV* mutants and the controls Bd21-3 and the outcrossed line *Bd NRPD1 (+/+)* are summarized as RdDM and Bd21-3, respectively. The following stresses were applied: c (control conditions), cold (2 °C on ice, 24h), drought (uprooting, 2:15 h), drug (chemical de-methylation with Zebularine (20 uM) and alpha-amanitin (2.5 mg/ml), 28 days), glyphosate (20 mM, four days), heat (42 °C, 8h), rice blast (*Magnaporthe oryzae* infection, four days), salt (300 mM NaCl, five days) and submergence (48 h). See material and methods for details.

### *HOPPLA* activity is increased in the *pol IV* mutants regardless of the stress applied

Fragmented eccDNAs or circles containing only one of the two LTRs (1-LTR circles) can be formed following reverse transcription by auto-integration, alternative end-joining in the virus-like particles [49,50] or by a recombination of the two LTRs of genomic copies [60,61]. Hence, 1-LTR circles do not necessarily imply LTR-RT mobility. In contrast, recent work indicates that 2-LTR circles are formed following the complete reverse transcription by non-homologous end-joining of an intact full-length linear RT copy that is capable of integrating into the genome [50]. Indeed, the detection of full-length 2-LTR circles of well characterized autonomous LTR-RTs such as *EVD* (*ATCOPIA93*) has been directly linked to their actual transposition [45]. As a complement to the assembly-based analysis, we thus followed a stringent approach to analyse our mobilome-seq data. We aligned reads to a library comprising artificial 3’LTR-5’LTR fusions of all full-length LTR-RTs annotated in the *B. distachyon* reference assembly [56]. This allowed us to specifically detect intact 2-LTR circles of extrachromosomal LTR-RTs capable of integrating into the genome. To control for possible traces of undigested genomic DNA that may also contain regions resembling LTR-LTR junctions and that may subsequently be amplified by the Phi29 enzyme during mobilome-seq [62], we also included publicly available genomic reads of all nine accessions in our analysis.

We found that several LTR-RTs formed eccDNA with 2-LTR junctions. Yet, most of them occurred sporadically and we did not observe a recurring stress-specific formation of 2-LTR circles for any of the 37 LTR-RT families with annotated full-length copies (**Fig 3A**). For instance, we found a very strong signal for RLC_BdisC031 that was solely detected in glyphosate-treated Bd21-3 plants and therefore not further considered in the analysis. In contrast, and in accordance with the assembly-based approach, the Angela element *HOPPLA* showed a recurrent formation of 2-LTR circles. However, this formation was not triggered by a specific stress and only occurred in the two independent *pol IV* mutants (**Fig 3B**). Finally, our attempt to transiently inhibit LTR-RT silencing using alpha-amanitin and Zebularine (a combination of inhibitors shown to increase the activity of LTR-RTs in *A. thaliana* and rice), did not result in the consistent activation of *HOPPLA* or other LTR-RTs in multiple accessions (**Fig 3A**).

**Fig 3.**
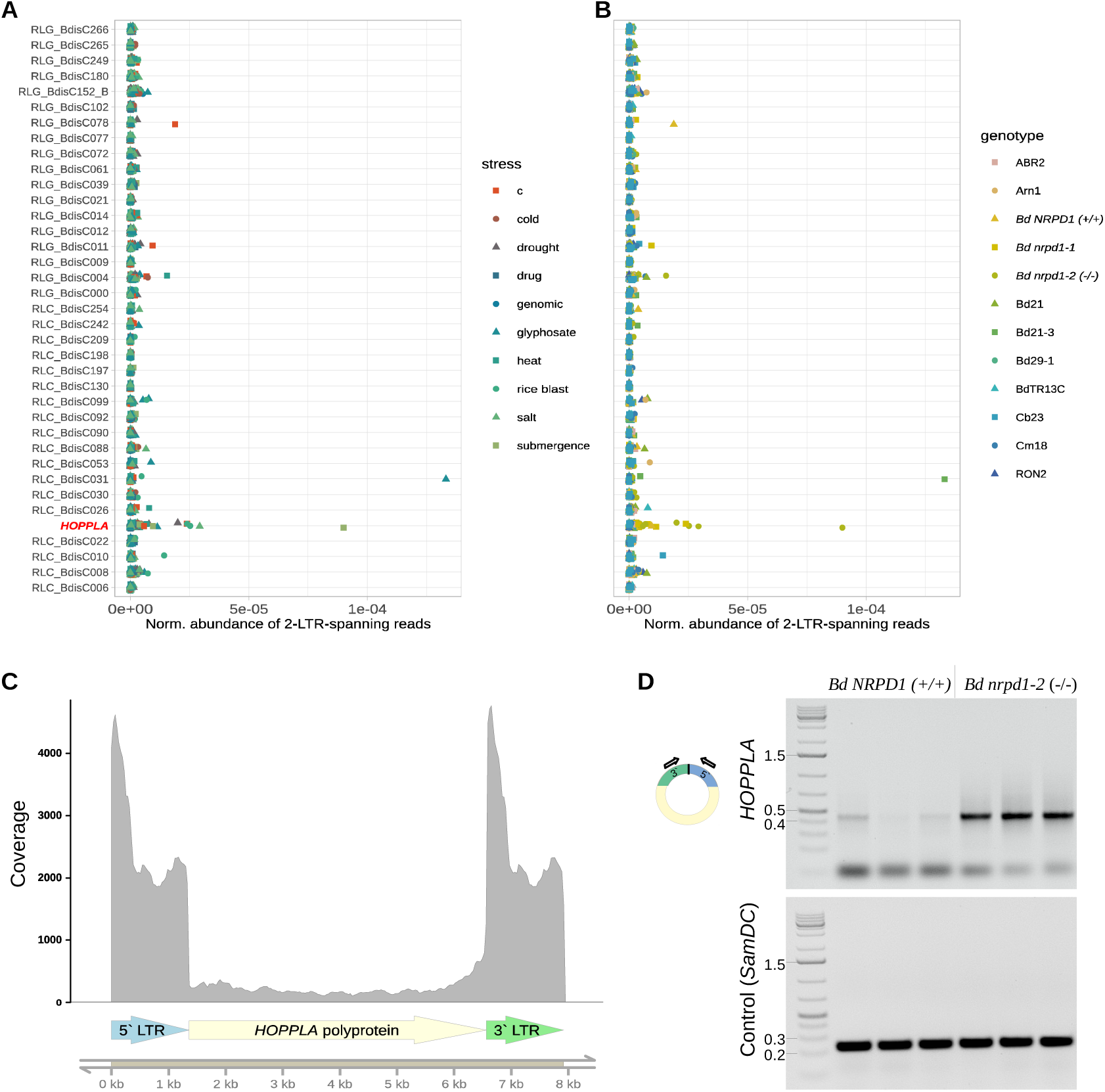
*HOPPLA* forms 2-LTR circles in the *pol IV* mutants. Normalized abundance of 2-LTR-junction spanning reads depending on the stress (**A**) and the genotype (**B**) of individual mobilome-seq samples. *HOPPLA* (RLC_BdisC024) is highlighted in red. The alignment of publicly available genomic reads (genomic) served as a control for the presence of genomic reads aligning to 2-LTR-junctions. See Fig 2 and methods section for details about the applied c (control) conditions and stresses. (**C**) Coverage of mobilome-seq reads of the *HOPPLA* consensus sequence of sample *Bd nrpd1-2 (-/-)* submergence stress. The structure of *HOPPLA* element is depicted. (**D**) Inverse PCR using total DNA not subjected to a rolling circle amplification for the confirmation of an increased amount of extrachromosomal 2-LTR circles of *HOPPLA* in the *Bd nrpd1-2* (-/-) mutant compared to the *Bd NRPD1 (+/+)* outcrossed line. Loaded PCR reactions with primers specific to the *HOPPLA* LTRs (top gel) and to the genomic control gene *SamDC* (Bradi5g14640) (bottom gel) are shown. Three biological replicates are depicted. Schematic representation of primer design for the inverse PCR is shown.

The mobility of *HOPPLA* in the *Bd nrpd1-2 (-/-)* was further confirmed by the fact that the alignment of mobilome-reads to the consensus sequence of *HOPPLA* resulted in a high coverage of the entire element (**Fig 3C**). Finally, the presence of 2-LTR circles of *HOPPLA* in *Bd nrpd1-2 (-/-)* was confirmed by an inverse PCR on total DNA that was not subjected to a rolling circle amplification, with outward facing primers specific to the two LTRs (**Fig 3D**). Notably, in the PCR, we also detected a faint signal for the outcrossed line *Bd NRPD1 (+/+)* suggesting a weak activity of *HOPPLA* in wild-type plants.

### Members of the *HOPPLA* family differ in activity

Because individual copies of the same LTR-RT family can differ in their activity [30], we also analysed the relative abundance of 2-LTR-spanning reads for each annotated full-length copy of *HOPPLA*. This analysis revealed a great diversity of eccDNA formation among individual copies of the *HOPPLA* family and confirmed the strongest activity of *HOPPLA* in the two *pol IV* mutants (**Fig 4A**). We also obtained a few reads spanning the 2-LTR-junction of the two most active *HOPPLA* copies (Bd3_22992889 and Bd4_25471847) in the *Bd NRPD1 (+/+)* control line (**Fig 4A**), which confirmed the weak but detectable band for the inverse 2-LTR PCR for *Bd NRPD1 (+/+)* (**Fig 3D**).

**Fig 4.**
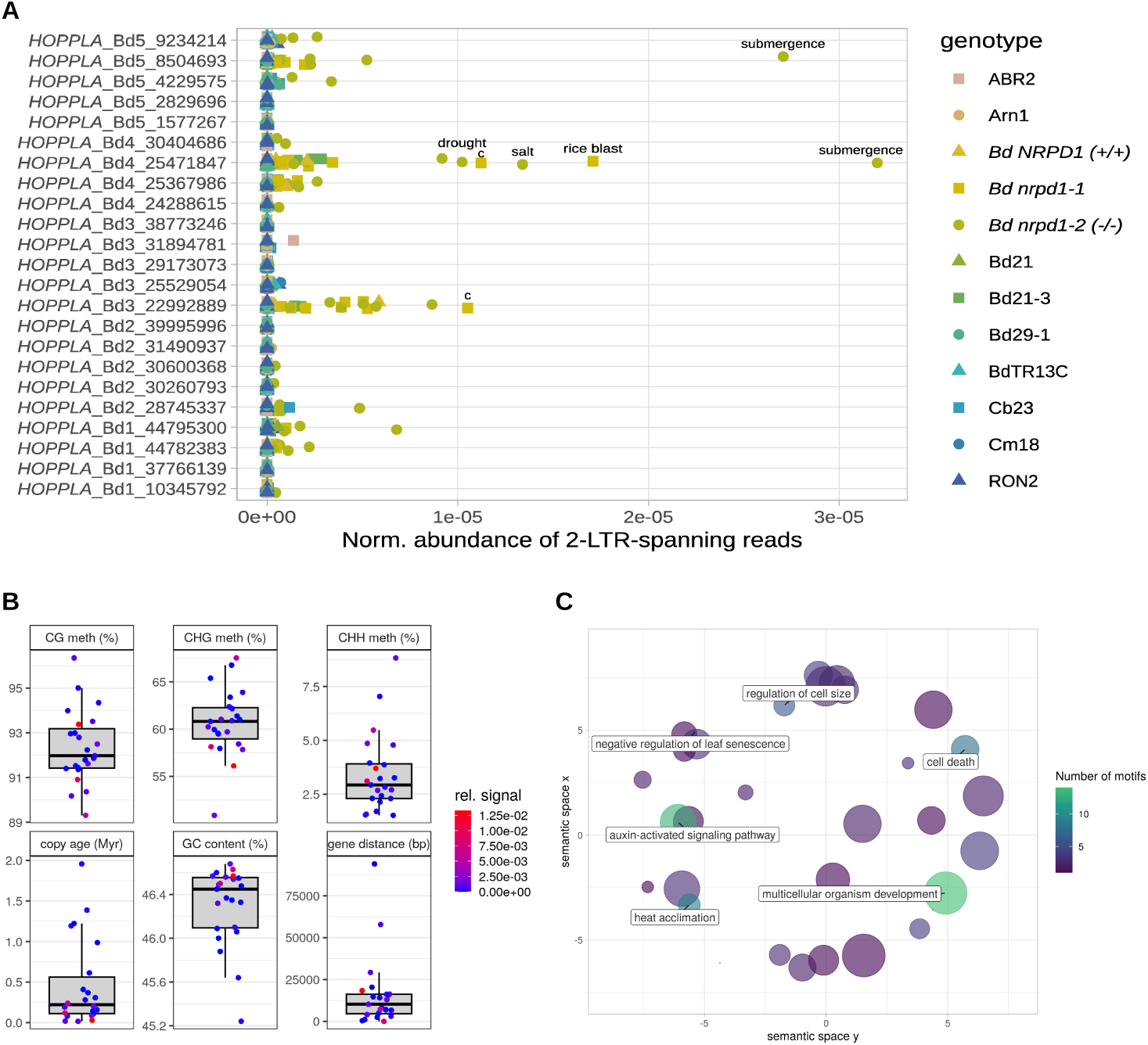
Members of the *HOPPLA* family differ in activity. (**A**) Normalized abundance of 2-LTR-junction spanning reads of individual full-length copies of the *HOPPLA* family. Samples with a high signal are labelled with c (control conditions) or the respective stress applied. See Fig 2 and material and methods for details. (**B**) Age (age_a), closest distance to gene (d_closest), GC content (GC), and methylation levels in CG, CHG and CHH contexts of all individual genomic full-length copies of *HOPPLA* in percent. The color indicates relative abundance of 2-LTR-junction spanning reads from the mobilome-seq in (**A)**. (**C**) GO enrichment analysis of transcription factors for which binding sites have been detected in the consensus sequence of *HOPPLA*. Colors indicate number of TF-binding sites found. GO-terms that occur at least six times are highlighted in the plot. All GO-terms and their number of occurrences are listed in S1 Table.

Using the meta information of individual *HOPPLA* copies described previously [56], we further assessed which genomic factors (DNA methylation, CG content, distance to the closest gene or age of the copy) were linked to the 2-LTR circle formation for individual *HOPPLA* copies. While no clear pattern emerged from this analysis, the most active copies of *HOPPLA* tend to be rather young (**Fig 4B**).

Since the stress- or tissue-dependent activity of LTR-RTs is mediated by the specific binding of transcription factor (TF), we screened the consensus sequence of *HOPPLA* for motifs of known TF binding sites. First, we validated this approach by analyzing one of most active copies (AT1G11265) of the heat-responsive *ONSEN* family of *A. thaliana*. As expected, a GO term analysis indicated a strong enrichment of heat-responsive TFs for this element (**S2 Fig**). In contrast to the well-known, stress-responsive *ONSEN* LTR-RT, the GO terms of TFs that could bind to *HOPPLA* may indicate that developmental processes and auxin-activated signaling pathways played a role in its activity, rather than specific stresses (**Fig 4C**).

### *HOPPLA* is targeted by Pol IV-dependent 24 nt siRNAs in the wild type and transposes in *pol IV* mutant plants

The pivotal role of Pol IV in producing TE-specific 24-nt siRNAs for RNA-directed DNA methylation has been demonstrated in many plant species including *A. thaliana* [27], rice [63] and for the Alesia family (RLC_BdisC010) in *B. distachyon* [64]. To confirm that the increased production of 2-LTR eccDNA circles of *HOPPLA* in mutants deficient in *B. distachyon* NRPD1 is correlated with a depletion of 24 nt siRNAs, we performed a small RNA blot including samples from the two *pol IV* mutants and their respective wild-type controls. Using a hybridization probe specific to the *HOPPLA* LTRs, 24 nt siRNAs were detected in the control lines Bd21-3 and *Bd NRPD1 (+/+)*, but not in either of the *pol IV* mutant lines (**Fig 5A**). This finding strongly suggests that *HOPPLA* is under control of the Pol IV-RdDM pathway, and that the absence of 24 nt siRNAs results in the upregulation and increased production of 2-LTR eccDNAs from *HOPPLA*. Furthermore, RNA-seq data from part of the same mutant panel shows that *HOPPLA* is the most upregulated LTR-RT family in the *Bd nrpd1-2 (-/-)* background compared to the *Bd NRPD1 (+/+)* control line, indicating that the reduction of 24 nt siRNAs is likely associated with an increased expression and subsequent formation of *HOPPLA* eccDNAs in both *polIV* mutants (**Fig 5B**).

**Fig 5.**
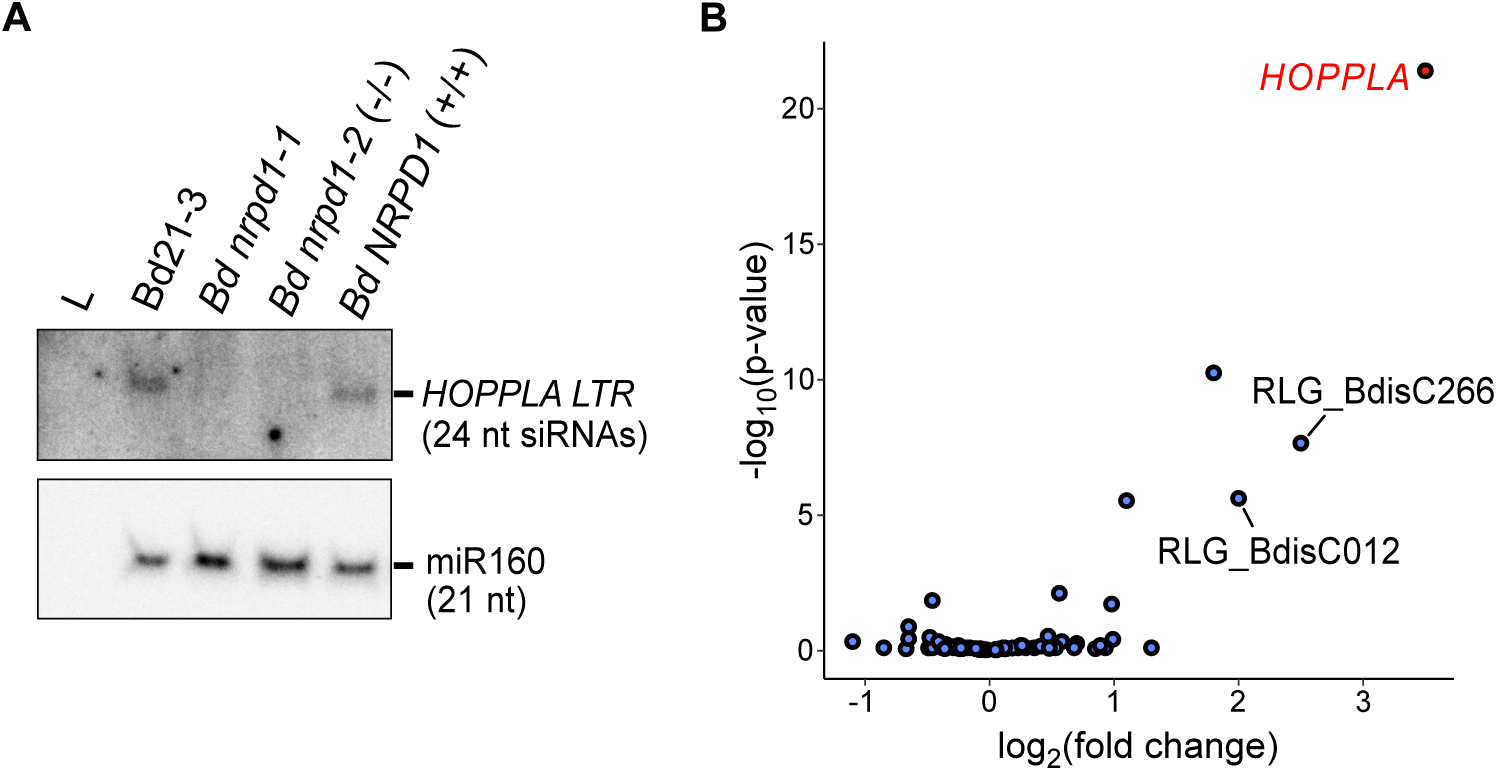
Loss of 24-nt siRNAs leads to an increased activity of *HOPPLA* in the Pol IV mutants. (**A**) Northern plot for the detection of 24-nt siRNAs specific to the 3′ LTR of *HOPPLA* in the *pol IV* mutants *Bd nrpd1-1* and *Bd nrpd1-2* (-/-) and their control lines Bd21-3 and the outcrossed line *Bd NRPD1* (+/+). (**B**) SalmonTE analysis of the expression of LTR-RTs in *Bd nrpd1-2* (-/-) relative to the outcrossed control line *Bd NRPD1* (+/+). LTR-RTs with a log2 fold change of at least two are labeled, *HOPPLA* is highlighted in red, three biological replicates were analysed.

To complete their life cycle, reverse transcribed extrachromosomal copies of LTR-RTs have to integrate into the host genome [41]. Because all our analyses congruently pointed to the activity of *HOPPLA* in the *Bd nrpd1-2 (-/-)* mutant, we sequenced the genome of seven individuals of *Bd nrpd1-2 (-/-)*, six *Bd NRPD1 (+/+)* plants and one wild-type Bd21-3 plant to detect new *HOPPLA* insertions. As TIPs were detected relative to the reference genome Bd21 (an accession closely related to Bd21-3 but not genetically identical), we first removed all conserved Bd21-3-specific TIPs detected in multiple lines. We manually curated all filtered candidate TIPs and showed that *HOPPLA* was the only family for which validated TIPs were identified in one of the re-sequenced *Bd nrpd1-2 (-/-)* plants (Bd1 38798495, Bd1 42205987, Bd4 28119639) (**S3A-S3C Fig**). This confirmed that the loss of Pol IV function led to an increased production of eccDNA, as well as actual transposition and accumulation of novel *HOPPLA* copies in the tested *Bd nrpd1-2 (-/-)* mutant. The presence of reads spanning the insertion site indicated that the detected *HOPPLA* insertions were heterozygous or probably somatic for the insertion Bd4 28119639, which exhibited a specifically low proportion of clipped reads. No TIPs were detected for any other LTR-RT family.

### Genome-wide association studies for pCN variations do not recover known components of RdDM

To decipher the genetic basis of *HOPPLA* accumulations in natural populations, we first performed a genome-wide association study (GWAS) using *HOPPLA* pCNs in the diversity panel of 320 natural accessions (**Fig 1B**) as a phenotype. We identified only one region with two significant peaks (FDR-adjusted p-value < 0.05, Bd5 6920000-6960000 and 7210000-7240000) obtained for the mobile *HOPPLA* family (**Fig 6**). Because inserted copies of *HOPPLA* may themselves lead to significantly associated regions in the GWAS as shown in *A. thaliana* [65], we first verified that there were neither TIPs [57] nor annotated reference insertions of *HOPPLA* in that region (**Fig 6**). As described above, our data suggested that the loss of 24-nt siRNAs in the Pol IV mutants was sufficient to mobilize *HOPPLA*. We therefore further tested whether any of the genes encoding subunits of Pol IV or Pol V would be localized in or near this region (window size 50 kb) (**S3 Table**). We did not detect any known Pol IV or Pol V-related genes, but instead found Bradi5g05225, an ortholog of the *A. thaliana* ROS1-associated methyl-DNA binding protein 1 (RMB1, AT1g63240) [66], to be co-localized with the peak.

**Fig 6.**
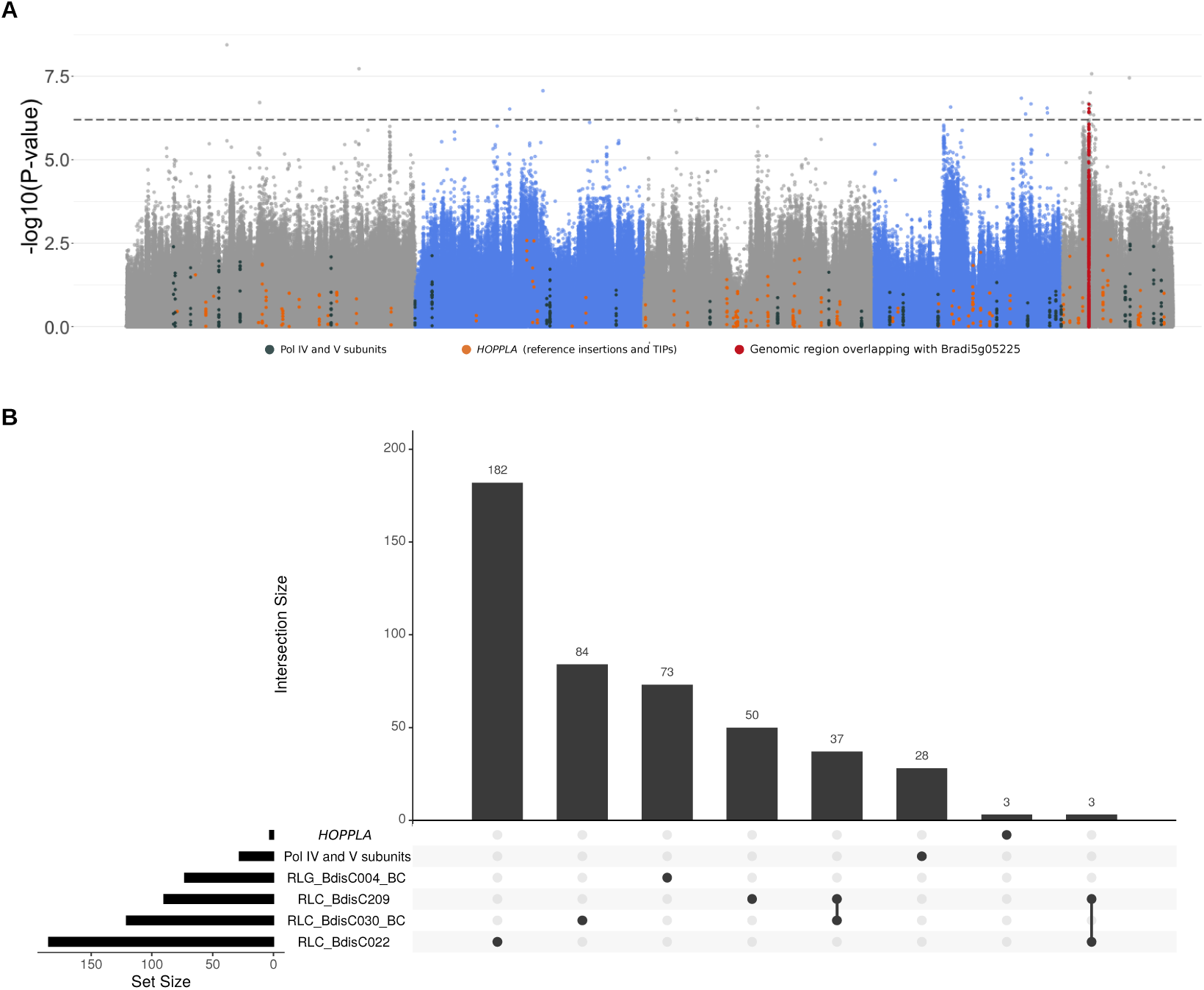
Genetic regions associated with pCN variations of recently active LTR-RTs. (**A**) Manhattan plot depicting the GWAS results of pCN variation of *HOPPLA* in 320 accessions of *B. distachyon.* Colored points indicate SNPs linked (+/- 10 kb) to known Pol IV and V subunits (dark grey), *HOPPLA* reference insertions and TIPs (orange) and the genomic region containing the candidate gene Bradi5g05225 (red). Threshold of *s*ignificance (false discovery rate adjusted p-value <0.05) is marked with dashed lines. A significant region containing the candidate gene Bradi5g05225 (window size 50 kb) is highlighted. (**B**) UpSet plot of genes in 20 kb windows surrounding significant regions with at least two SNPs above the threshold of significance (FDR-adjusted p-value <0.05 for *HOPPLA*, RLC_BdisC209 and RLC_BdisC022 and Bonferroni correction for RLC_BdisC030 and RLG_BdisC004) of the five most recently active LTR-RT families in *B. distachyon*. To visualize potential overlaps, a list of the components of the Pol IV and Pol V holoenzymes is included in the UpSet plot.

To test whether genomic regions might be recurrently associated with their pCNs, we finally extended the GWAS analyses to the four other most recently active families (RLC_BdisC022, RLC_BdisC030, RLC_BdisC209 and RLG_BdisC004) (**S4 Fig**). We extracted the candidate genes for each of the five families (see **S3 Table** and Material and Methods). Apart from the two closely related families RLC_BdisC030 and RLC_BdisC209 that shared the majority of their GWAS candidates, and RLC_BdisC022 and RLC_BdisC209 that shared three genes, we found no overlap of annotated loci potentially contributing to the pCN variations of recently active families (**Fig 6**).

## Discussion

Understanding the dynamics of TEs and their role in adaptation is currently one of the major challenges in the field of evolutionary genomics. The fact that mobile TEs are a source of epi/genetic diversity and potential drivers of evolution has been demonstrated in many organisms including fungi [67], insects [68], mammals [69] and plants [70]. However, while there are a number of examples showing that certain TE insertions facilitated the adaptation to changing environments (for review [71]), TEs are generally harmful and therefore controlled by complex silencing mechanisms. To foster our understanding of TE activity, we investigated the environmental conditions and genetic factors associated with the accumulation and mobility of LTR-RTs in plant genomes. By measuring LTR-RT pCNs in a panel of 320 *B. distachyon* natural accessions, we show that the intra-specific variations of pCNs of RLC elements, but not the pCNs of the generally older and more abundant RLG elements [56], separate accessions according to their genetic cluster of origin. This is even more striking for members of the Angela (RLC_BdisC022, RLC_BdisC024, RLC_BdisC030, RLC_BdisC209) and the Tekay (RLG_BdisC004) family, which are the youngest families in *B. distachyon* [56]. Highly polymorphic among natural accessions of *B. distachyon* [17,56], they are expectedly the main drivers of lineage-specific expansions of pCNs in our study system. Hence, we do not only confirm that LTR-RT families in *B. distachyon* globally differ in size [56] but also demonstrate that the accumulation of genomic sequences derived from specific families varies significantly among natural accessions.

The transcriptional activity of LTR-RTs can be triggered by specific environmental stresses [29,34]. Given that *B. distachyon* occurs in a wide range of different habitats in the Mediterranean area [53], this characteristic feature of LTR-RTs provides a potential explanation for the pCN variation we observed across natural accessions [18,65]. Yet, for the large majority of LTR-RT families, pCNs correlate only moderately with environmental factors. Consequently, our genomic data do not support a large effect of the environment on LTR-RT activity in *B. distachyon*. While this result could seem startling, it is not completely surprising. Indeed, many LTR-RT families, and especially the old RLG elements, do not show signs of increased activity in the recent past in *B. distachyon* [56]. Considering that their copy number expansions took place in a climate that has drastically changed following the last glacial maximum [53], a limited link between their activity and the current environmental conditions is actually expected for most families. In contrast, the lack of correlation between current bioclimatic variables and copy number variation for families with ongoing activity (RLC_BdisC022, RLC_BdisC024, RLC_BdisC030, RLC_BdisC209 and RLG_BdisC004; [56]), suggests a more complex mechanism than their pure dependence on specific stresses in certain environments. This hypothesis is supported by previous findings in *A. thaliana*. Indeed, a minor impact of the environment on transpositional activity was also found in this species, where the two most associated environmental variables (‘seasonality of precipitation’ and ‘diurnal temperature range’) only explained about 9 % of the observed variation [18].

Genetic factors are well-known to be essential in regulating LTR-RT activity [72–75]. As the loss of main players of the RdDM silencing pathway leads to increased TE activity [18,27,33,63], the two *B. distachyon* Pol IV (NRPD1) mutants provided an ideal functional tool to experimentally validate our *in silico* analysis. Since transcriptionally active LTR-RTs are not necessarily able to transpose [76], we used a mobilome-seq approach to detect TE-derived eccDNAs and transpositionally active LTR-RT families. We deliberately followed a very stringent approach for analysing the data and by doing so, identified *HOPPLA* as the only highly active LTR-RT family in *B. distachyon.* Indeed, *HOPPLA* is the only family for which we further detect newly inserted copies in the Pol IV mutant.

The non-stress-specific activity of *HOPPLA* in the two independent Pol IV mutants and limited activity of other elements supported our *in silico* approach and strengthened the idea that genetic, rather than environmental stresses, are major drivers of LTR-RT activity in *B. distachyon*. While we cannot exclude the possibility that other specific stress conditions may trigger the eccDNA formation of other young autonomous LTR-RTs, specifically for *HOPPLA*, these results are also in line with our TF binding sites analysis. In contrast to the heat-responsive *A. thaliana* element *ONSEN*, for which we predominantly recovered TF-binding sites associated with heat response, *HOPPLA* seemed to be targeted by TFs involved in developmental processes and auxin signaling [77]. As this study already covers a broad range of (a)biotic stresses, follow-up studies should therefore address the question of whether the activity of *HOPPLA* or other families differs between tissues or developmental stages, as also observed for the endosperm-specific mobility of *PopRice* in rice, for example [45]. Strikingly, despite the central role of Pol IV in the RdDM pathway, we did not observe bursts of multiple LTR-RT families but instead found that the loss of 24 nt siRNAs specifically activated individual copies of the *HOPPLA* family. Interestingly, we also detected a weak signal for 2-LTR eccDNAs in the Bd21-3 wt and the outcrossed line *Bd NRPD1 (+/+)* but not in other natural accessions. This suggests that the accession-specific composition of the mobilome, and hence the genetic background of the *pol IV* mutant line, plays an important role in LTR-RT activity. Related to this, we sporadically observed very strong signals for individual samples, which could indicate an accession-specific response of the mobilome to certain triggers.

Given that pCNs vary greatly among genetic clades, assessing the effect of a genetic mutation of major components of the RdDM pathway in a set of genetically diverse natural accessions would be timely, yet labor-intensive as transformation works more efficiently in the Bd21-3 background than in the other accessions tested. Our attempt to transiently reduce LTR-RT silencing in multiple accessions from different genetic clades using the chemical inhibition of Pol II and DNA methyltransferases [52] did not result in an increased activity of *HOPPLA* or members of any other LTR-RT family. In addition, and despite the differences of activities observed among individual *HOPPLA* copies, we could not detect, in the present study, a clear link between their activity and GC contents or methylation states. Taken together, these findings suggest that the specific function of the canonical RdDM with Pol IV, rather than generic DNA methylation states are regulating *HOPPLA* activity. Yet, our GWAS failed to recover major components of the RdDM pathway. Instead, the diversity of activity within the *HOPPLA* family may suggest that the presence of single active copies could determine the fate of an entire family. In addition, pCNs are also dependent on the removal rate of LTR-RT families, resulting in the formation of TE-fragments and soloLTRs, which varies greatly in *B. distachyon* [56]. This complexity of parameters affecting the dynamics of LTR-RT might explain why none of the genes known to be involved in silencing LTR-RTs are associated with the pCN variation of *HOPPLA* or any recently active family. Our candidate locus containing Bradi5g05225, a gene related to RMB1 whose loss of function has been shown to result in DNA hypermethylation [78], remains nonetheless a great candidate for functional validation.

Altogether, our work confirms that LTR-RTs in *B. distachyon* are ‘well-behaved’ [17] and that the evolutionary consequences of their mobility are hard to study in real-time. Indeed, while mobilome-seq revealed a sporadic activity for other families, we only found recurring activity and new insertions of *HOPPLA* in the *pol IV* mutant. These results somewhat contrast with our previous population genomics analyses which clearly indicate an ongoing activity of several LTR-RT families in natural accessions. We propose that the activity of LTR-RTs is relatively low and might depend on a complex interaction between genetic factors, developmental stages and, more marginally, the punctual occurrence of stresses. This study not only elucidates fundamental mechanisms of LTR-RT-dynamics of an undomesticated grass in the wild, but may also be relevant to better understand the biology of mobile elements in more complex genomes.

## Materials and methods

### Estimation of LTR-RT pCNs

We used publicly available genomic reads of 320 sequenced natural accessions of *B. distachyon* [53,58,59,79,80] to assess the natural variation of copy numbers of LTR-RTs. Because sequencing depth differed substantially between accessions [53], we first downsampled all fastq files to the read number of the sample with the lowest number of reads (4.230.721 reads) using the reformat.sh function of BBtools (v 38.75, BBMap Bushnell B., sourceforge.net/projects/bbmap/). Downsampled reads were aligned to the TREP consensus sequences of LTR-RTs and the reference assembly of Bd21 (v 3.0) using BWA-MEM (v 0.7.17-r1188) [81] with the -M and the -a options set, hence outputting all alignments found. Coverages of LTR-RTs and the reference assembly were assessed using bedtools (v 2.30.0) [82] genomecov with the -d and -split parameters set. The assessment of the global coverage of the reference genome allowed us to compensate for potential differences in read length and/or quality. A proxy for copy numbers was thus obtained by normalizing the coverage signals of each of the LTR-RT consensus sequences by the coverage of the entire reference assembly and by correcting for the length of the consensus sequences. pCN raw data were processed using R (v 3.6.3 and 4.0.2) [83] in Rstudio [84].

Variation in pCNs across the 320 natural accessions was visualized with a heatmap drawn with the heatmap() function natively provided in R version 4.0.2. We computed pairwise genetic distances between accessions with the R package pvclust v 2.2.0 [85]. The resulting tree was used to order accessions phylogenetically on the heatmap. PCAs based on pCNs were obtained with the R package ggbiplot v 0.55 [86].

To test for an association between pCNs and environmental variables, we retrieved information about climatic variables at each local site from [53]. Linear mixed model analyses where the pCN per LTR-RT family was entered as the response variable, the bioclimatic variables entered separately as fixed factors and the clade of origin as random factors to account for population structure were ran with the R package lme4 [87]. The part of the variance explained by the fixed-(marginal R^2^) were computed following [88] and visualized as bubble plot with the R package ggplot2 [89]. Classical linear models were run in base R.

### Plant material, growth conditions and stresses for mobilome-seq

*Brachypodium distachyon* natural accessions used in this study comprised Bd21, Bd21-3, Cm18, Cb23, ABR2, Bd29-1 BdTR13c, RON2 and Arn1. Because Pol IV is known to play an important role in LTR-RT silencing in plants [27,90,91], we also included the sodium azide mutagenized *pol IV* mutant line NaN74 (*Bd nrpd1-1*) [64,92], the T-DNA insertion *pol IV* mutant line JJJ18557 Nr31 [93] *Bd nrpd1-2 (-/-)* and a corresponding sibling, outcrossed control line *Bd NRPD1* (+/+) in the background of the natural accession Bd21-3. For *in vitro* experiments, seeds were soaked for 4 h in tap water and, without damaging the embryo, the lemma was carefully peeled off. Seeds were then surface-sterilized for 30 seconds in 100% ethanol and immediately rinsed three times with sterile tap water. Surface-sterilized seeds were placed with the embryo facing down and at an angle of about 30° towards the side, onto solid ½ MS-medium (2.15 g/L MS basal salt without vitamins (Duchefa Biochemie, Haarlem, NL)), 0.5 g/L MES-Monohydrate, 10 g/L sucrose, pH 5.8 (KOH), 0.25 % Phytagel (Sigma-Aldrich, St. Louis, USA) in ‘De Wit’ culture tubes (Duchefa Biochemie, Haarlem, NL). Plants were grown at 24 °C (day) / 22 °C (night), 16 h light under controlled conditions in an Aralab 600 growth chamber (Rio de Mouro, PT) for 25 to 29 days until the onset of stresses. For salt stress, seedlings were transplanted to solid ½ MS-medium supplied with 300 mM NaCl and grown for five days at 24/22 °C, 16h light. A solution of sterile-filtrated Glyphosate (Sintagro AG, Härkingen,CH) (20 mM, diluted in water) was applied to leaves using a piece of soaked sterile filter paper and plants were incubated for four days at 24/22 °C, 16h light. Drought stress was induced by uprooting plants from the medium and incubating them for 2:15 h at 24 °C in the light. Before sampling, plants were allowed to recover for two hours on fresh ½ MS-medium. For the infection with *Magnaporthe oryzae* (rice blast) six isolates (FR13, Mo15-27, 9475-1-3, IK81, M64 and Mo15-19) with spore concentrations between 130‘000-200’000 K spores per isolate per mL sterile water, supplied with 0.2 % Tween 20 were mixed and applied with a cotton swab to plant leaves. Plants were incubated for 24 h in the dark (24/22 °C) and then grown for another three days at 24/22 °C, 16h light. For heat stress, plants were incubated for 8 h at 42 °C. Before sampling, heat-stressed plants were allowed to recover for 16 h at 24/22 °C. Cold stress was induced by incubating plants for 24 h at 2 °C on ice at 16 h light. Prior to sampling, plants were allowed to recover for two hours at 24 °C in the light. For submergence stress, two small holes were drilled just above the growth medium and at the top through the wall of the culture tubes. Tubes were then inverted and submerged upside down for 48 h at 24/22 °C, 16h light using a custom rack in a plastic beaker filled with 2.5 liters of 24 °C tap water. In this way, it is possible to submerge plant leaves without the medium coming into contact with the water. Chemical de-methylation of DNA was conducted according to [52] by germinating and growing plants for 28 days on ½ MS-medium supplied with a mixture of Zebularine (Sigma-Aldrich, St. Louis, USA) and alpha-amanitin (Sigma-Aldrich, St. Louis, USA). Because the drug treatment severely affected the growth of seedlings, we omitted a treatment of mutant plants and used reduced concentrations of 20 uM (Zebularine) and 2.5 mg/ml (alpha-amanitin), respectively for all natural accessions.

### Mobilome sequencing and validation of eccDNAs

DNA was extracted using the DNeasy plant kit (Qiagen, Venlo, Netherlands) according to the protocol of the manufacturer. DNA concentration was measured using the Qubit high sensitivity kit (Invitrogen, Waltham, USA). Mobilome sequencing was performed according to [45] using pooled DNA of two biological replicates per sample. For this, 50 ng of DNA from both biological replicates were pooled and diluted to a volume of 58 μL. To enrich eccDNA, DNA was first purified using the GENECLEAN kit (MP Biomedicals, Santa Ana, USA) according to manufactures recommendations using 5 μL glass milk with an elution volume of 35 μL. Thirty μL of the eluate were digested using the Plasmid-Safe ATP-dependent DNase (Biosearch Technologies, Hoddesdon, UK) for 17 h at 37 °C. The digestion product was then subjected to an ethanolic precipitation and the precipitated eccDNA amplified using the illustraTempliPhi Amplification Kit (Cytiva, Marlborough, USA) according to [45] with an extended incubation time of 65 h at 28 °C. The templiphi product was diluted 1:10, quantified using the Qubit high sensitivity kit and 120 ng per sample were used for library preparation. Sequencing libraries were prepared using the Nextera DNA Flex Library Prep and the Nextera DNA CD Indexes (Illumina, San Diego, USA). Quality of libraries were assessed using the Tape Station (Agilent Technologies, Santa Clara, USA) with High Sensitivity D1000 screen tapes and concentrations were measured using the Qubit high sensitivity kit. Up to 12 indexed libraries were pooled and sequenced with an Illumina MiSeq sequencer using the MiSeq reagent kit v3 (600 cycles) (S6 **Table)**. Raw reads have been uploaded to ENA (accession number PRJEB58186).

The presence of extrachromosomal circular copies of *HOPPLA* (RLC_BdisC024) was validated by an inverse PCR using 7 ng/μL total DNA. Input quantities of DNA were controlled using primers specific to the S-adenosylmethionine decarboxylase (SamDC) gene (Bradi5g14640) that have previously been shown to be efficient and are therefore also used for the control reaction amplifying reference genes in real-time PCRs in *B. distachyon* [94]. Sequences of primers are listed in **S5 Table**.

### Analysis of Mobilome-seq

Reads were trimmed using the BBDuk tool of BBtools (BBMap (v 38.75, Bushnell B., <u>sourceforge.net/projects/bbmap/)</u> with the parameters qtrim = rl and trimq = 20. Reads originating from organelles were removed by aligning reads to the chloroplast genome (NC_011032.1) [95] and the mitochondrion genome (v 1.0.0) of *B. distachyon* using BWA-MEM (v 0.7.17-r1188) [81] with the - M parameter set. Unmapped reads were isolated using samtools (v 1.13) [96] view -b -f 4 and bedtools (v 2.30.0) [82] bamtofastq.

Organelle-filtered mobilome reads were assembled using the SPAdes genome assembler (v 3.13.0) [97]. From each assembly, the top ten contigs were extracted and jointly aligned to the reference assembly of Bd21 (v 3.0) using BWA-MEM (v 0.7.17-r1188) with the -M parameter set. Bam files were converted into bed files using bedtools (v 2.30.0) bamtobed with the -split option set and overlapping contigs were merged using bedtools merge with the -o distinct, count, count_distinct and -c 4 parameters set. Assembled, circle-forming regions were annotated with bedtools intersect using the version 3.1 annotation of the reference assembly and the annotation of all full-length LTR-RTs [56] of the reference assembly. Annotated regions were extracted with bedtools getfasta and all sequences longer than 2 kb were isolated using SeqKit seq (v 0.11.0) [98]. Circle-forming regions that occurred in less than three samples were not included in the analysis.

To specifically detect mobilized LTR-RTs, we first extracted all annotated full-length LTR-RTs of the Bd21 reference assembly [56]. Using a custom python script, we then merged the last 300 bp of the 3’ to the first 300 bp of the 5’ LTR to obtain a ‘tail-to-head’ library containing all annotated full-length LTR-RT copies annotated in Bd21. We then aligned organelle-filtered mobilome reads to the tail-to-head library of LTR-RTs and used bedtools (v 2.30.0) intersect to extract aligned reads that were spanning the 2-LTR junction and that aligned to at least 5 bp of both LTRs. The coverage of the junction-spanning reads was calculated using deeptools (v 3.5.1) [99] with the parameters -bs 1, --ignoreDuplicates --outRawCounts set. To account for differences in sequencing depth, the obtained coverage for 2-LTR-junction spanning reads was normalized with the total coverage obtained with bedtools (v 2.30.0) genomecov with the - d and -split parameters set, from the alignments of filtered reads to the reference assembly of Bd21 (v 3.0) obtained from Phytozome 12 [55] generated by BWA-MEM (v 0.7.17-r1188) [81]. To control for potential traces with undigested genomic DNA that may contain inserted LTR-LTR junctions, we also aligned publicly available reads for each of the accessions, and measured their relative abundance as described above. To plot the overall activity per family (**Fig 3A and 3B**), normalized signals were summed up for every individual TE family. Otherwise (**Fig 4A**) signals were plotted for each individual *HOPPLA* full-length copy.

To visualize the coverage of the *HOPPLA* TREP consensus sequence, reads were aligned using BWA-MEM with the -M and -a options set. Aligned reads were visualized using the packages GVIZ (v.1.28.3) [100] and RTRACKLAYER (v.1.44.4) [101] in R and Rstudio.

### mRNA-sequencing and small RNA northern blotting

Leaves of 4-week-old *B. distachyon* plants were ground in liquid nitrogen and 500 μL of this powder was subjected to TRIzol extraction following the supplier instructions (Invitrogen, CA, USA). 20 µg of total RNA was treated with DNase I for 30 min., then repurified via phenol-chloroform extraction and ethanol precipitation. DNase-treated total RNA samples were sent to Fasteris/Genesupport (Plan-les-Ouates, Switzerland), subjected to poly(A)-tail selection, and then aliquoted for library construction via the Illumina TruSeq Stranded mRNA Library Prep kit. Resulting stranded polyA+ RNA-seq (mRNA-seq) libraries were sequenced on an Illumina NovaSeq 6000. The raw paired-end read data were deposited at the NCBI Gene Expression Omnibus (GEO accession: GSE243693).

For the small RNA blot analysis, 200 µg of each total RNA were size-fractionated using the RNeasy Midi Kit (QIAGEN), as described previously [64]. Low molecular weight (LMW, <200 nt) RNAs are not bound by the silica membrane of the columns and were isolated from the collected flow-through and wash aliquots. LMW RNAs were precipitated overnight using isopropanol. Following a centrifugation step (45 min. at 24000 x g, 4°C) and the removal of the supernatant, the pellet was washed with 75% ethanol, centrifuged (15 min. at 24000 x g, 4°C), dried at RT for 20 min. then at 65°C for 5 min., and resuspended in 41 µL of DEPC-treated MilliQ water. LMW RNAs were quantified using a Nanodrop device and 12.3 µg of LMW RNAs from each sample were loaded into the 16% polyacrylamide gel [64]. After running, transfer and UV crosslinking, membrane was prehybridized in PerfectHyb Plus buffer (Merck, Darmstadt, Germany) at 35°C and then and hybridized at 35°C with the Klenow internally-labeled probe (*HOPPLA*), or with the 5’-end labeled probe (miR160) [64]. After overnight hybridization, washing was performed at 37°C. Signal detection requires 5-7 days exposure for *HOPPLA* and 1-2 days for miR160. Oligonucleotide sequences for the probes are listed in **S5 Table.**

### LTR-RT expression analysis

RNA-seq raw reads of *Bd nrpd1-2 (-/-)* and *Bd NRPD1 (+/+)* were trimmed for adapters using fastp (v 0.23.2) [102] with the following options: --qualified_quality_phred 15 --unqualified_percent_limit 4 -- n_base_limit 20 --low_complexity_filter --overrepresentation_analysis --correction -- detect_adapter_for_pe. Cleaned reads were then analysed using SalmonTE (v 0.4) [103] to measure global expression of LTR-RTs. LTR-RT consensus sequences of *B. distachyon* obtained from the TRansposable Elements Platform (TREP, https://trep-db.uzh.ch/) were used to generate the custom library for SalmonTE. Default options of SalmonTE quant and test function were used to quantify expression and to perform statistical analysis. Expression data were plotted using R (v 3.6.3) in RStudio (v 7d165dcf).

### Motif analysis

The consensus sequence of *HOPPLA* was screened for known transcription factor binding sites obtained from the PlantTFDB [104] using FIMO (v 5.1.1) [105]. To functionally annotate transcription factors that could bind to *HOPPLA,* we used GO-terms of the Gramene (release 50) database [106] downloaded from the platform agriGO (v 2.0) [107]. Generic, TF specific GO terms (GO:0003700, GO:0006355, GO:0005634, GO:0003677, GO:0043565, GO:0046983, GO:0003682 GO:0045893) such as ‘positive regulation of DNA-templated transcription’ were removed from the list of GO terms as they would interfere with the downstream analysis. The remaining GO terms of transcription factors potentially binding to *HOPPLA* were visualized with REVIGO [108] using the ‘SimRel’ semantic similarity measure, the option ‘small’ and the GO terms of the *Oryza sativa* Japonica Group. The total number of occurrences of individual GO terms was taken into account with the option ‘higher value is better’. GO terms occurring more than five times were labelled in the plots. As a proof of concept, we followed the exact same approach using the sequence of one of the most active *ONSEN* copies (AT1G11265) and the GO terms of *Arabidopsis thaliana*.

### Detection of novel *HOPPLA* insertions in *Bd nrpd1-2 (-/-)*

DNA of adult plants was extracted using the DNeasy plant kit (Qiagen, Venlo, Netherlands) according to the protocol of the manufacturer subjected to whole genome sequencing. Reads were trimmed using the BBDuk tool of BBtools (BBMap (v 38.75, Bushnell B., <u>sourceforge.net/projects/bbmap/</u>) with the parameters qtrim = rl and trimq = 20. Trimmed reads were aligned to the reference assembly of Bd21 (v 3.0) using BWA-MEM (v 0.7.17-r1188) [81] with the -M parameter set. Samtools (v 1.13) [96] was used to obtain sorted and indexed bam files. TIPs were detected with detettore (v 2.0.3) (https://github.com/cstritt/detettore) with the options –require_split, -q 30 and using the consensus sequences of LTR-RTs of *B. distachyon* (TREP-database) and the annotation of all full-length LTR-RTs of Bd21 [56]. Because both *Bd nrpd1-2* lines were in the Bd21-3 background we were able to exclude all Bd21-3 specific TIPs by removing those insertions that were detected in multiple individuals with more than one genetic background. Remaining TIPs were manually curated using the genome browser IGV (v 2.15.4.12) . *HOPPLA* TIPs were visualized with JBrowse 2 (v 2.6.1) [109]. Raw genomic reads of the re-sequencing of *Bd nrpd1-2 (-/-), Bd NRPD1 (+/+)* and Bd21-3 have been uploaded to ENA (accession number PRJEB73379).

### GWAS for pCNs

GEMMA 0.98.5 [110] was used to test for associations between SNPs [53] and the LTR-RT families pCNs, while correcting for population structure [53,58]. Α centered relatedness matrix was first created with the option -gk 1 and association tests were performed using the option -maf 0.05 to exclude rare alleles, and the default SNP missingness threshold applied by GEMMA that excludes SNPs with missing data in more than 5% of the accessions. We selected 20 kb genomic regions with a 10 kb overlap that contained at least two SNPs above the False Discovery Rate of 0.05 or Bonferroni correction threshold as candidate region using the R package rehh (v 3.2.2) [111]. Genes overlapping with candidate regions were selected with the BEDTOOLS (v 2.26.0) [82] intersect command using the version 3.1 of the *B. distachyon* annotation file (https://phytozome-next.jgi.doe.gov). The UpSetR [112] R package was used to visualize the intersections of significant genes between the variables. Protein constituents of the Pol IV and Pol V enzymes (see **S4 Table**) were downloaded from the plant RNA polymerase database http://rna.polymerase.eu/.

## Acknowledgements

We thank the Genetic Diversity Center Zürich for providing the infrastructure for the mobilome-seq. This manuscript was professionally edited by Dr Emmanuelle Botté, from Manuscribe (https://manuscribe.com.au/).

## Author contributions

M.T. and A.C.R. conceived the study and planned experiments. M.T., B.K., K.R., C.H., B.R. and M.B. generated data. M.T., A.C.R., N.M. and W.X. analysed data. C.M. handled sequencing data. D.L., J.V. and R.S. provided mutagenized germplasm and online TILLING/T-DNA browsing tools. C.H., B.R. and T.B. identified and screened for the *pol IV* null mutants. M.T and A.C.R wrote the paper with contributions from T.B., C.S. All authors read the manuscript.

## Supporting Information - Figures

**S1 Fig.**
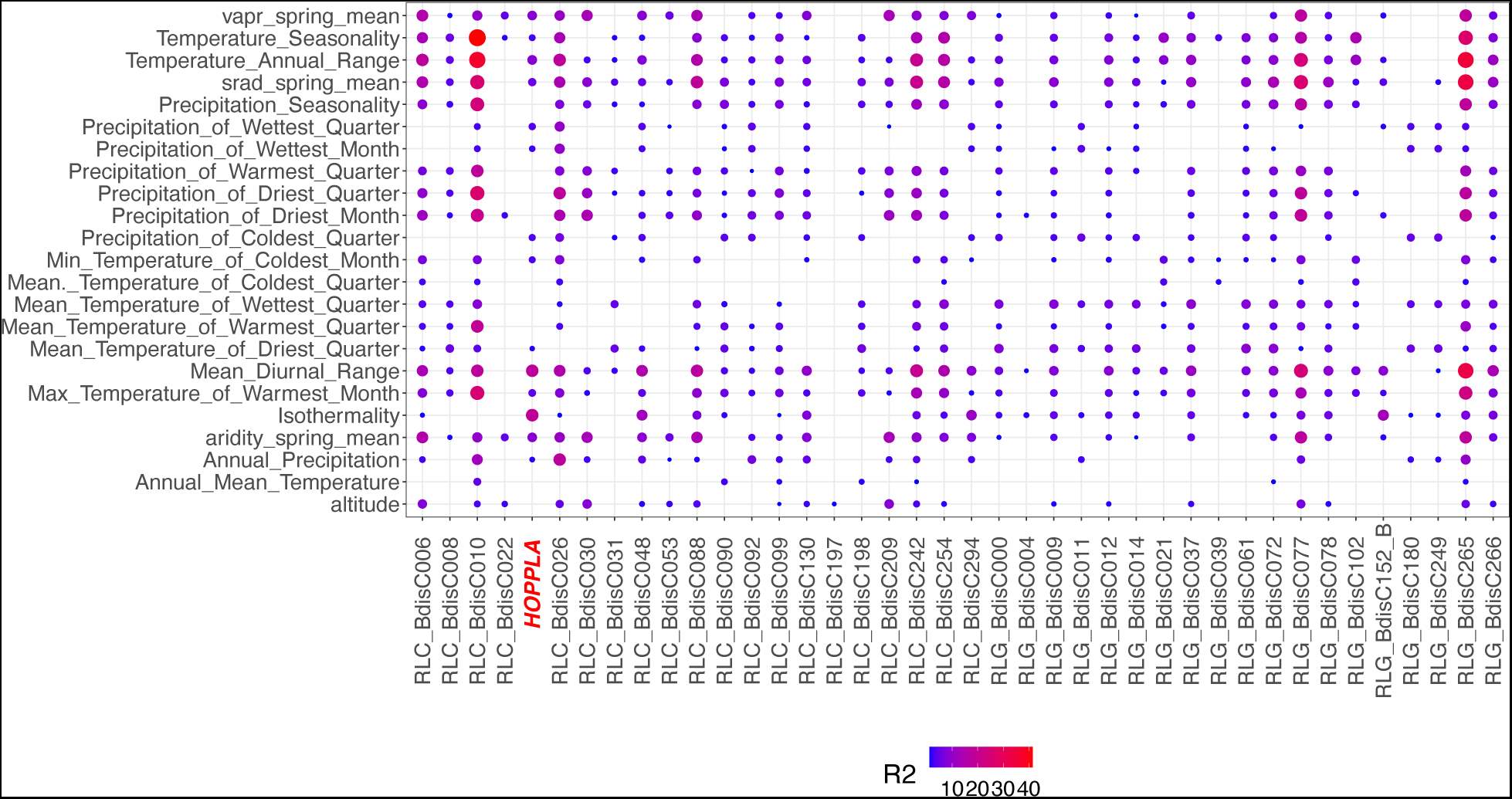
Correlation of bioclimatic variables with pCNs when not correcting for population structure. Colors and sizes of bubbles show the part of the variance (R^2^) explained by the bioclimatic variables in %

**S2 Fig.**
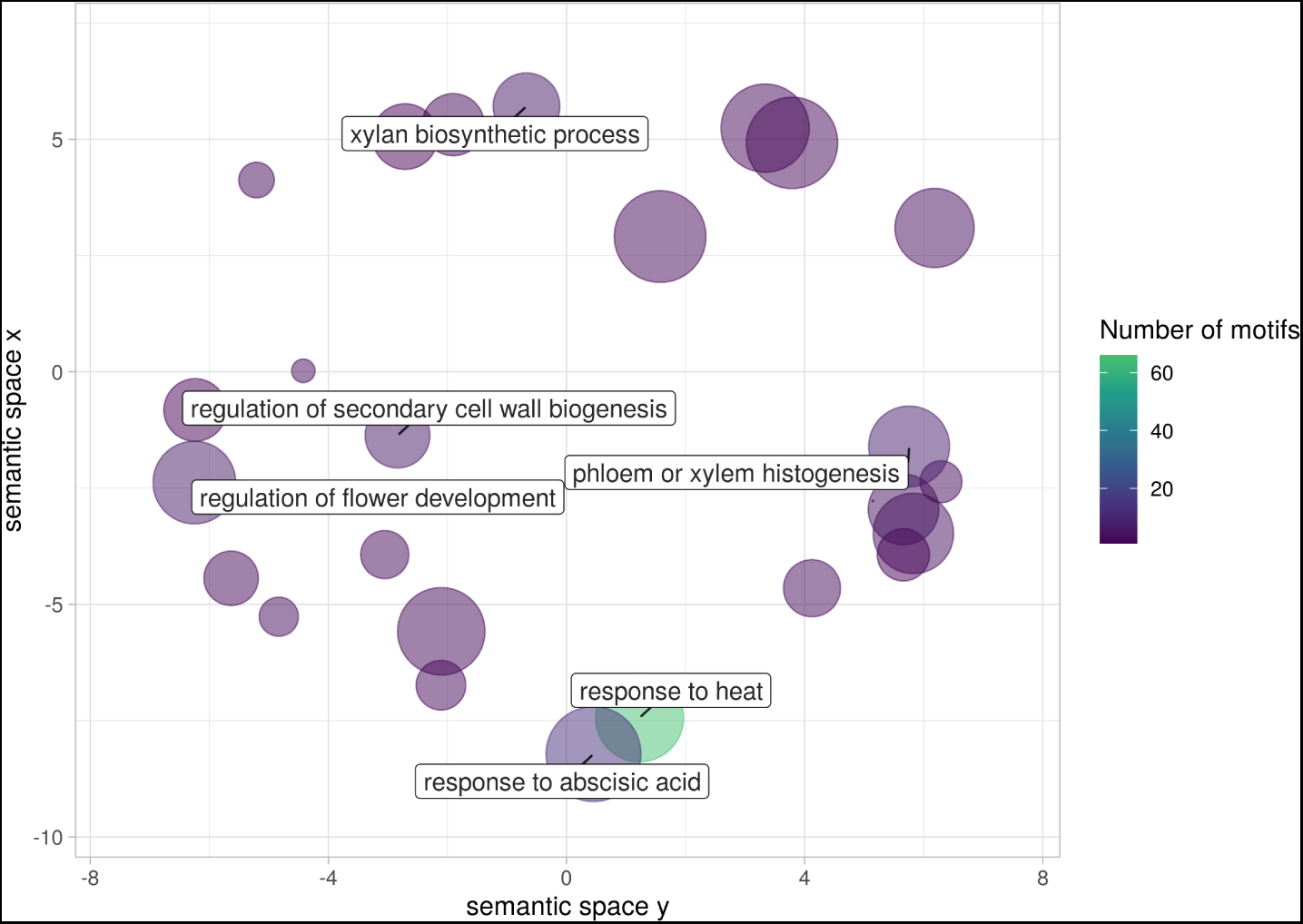
TFs binding to *ONSEN* are heat-stress inducible. GO-enrichment analysis of transcription factors for which binding sites have been detected in AT1G11265, a member of the heat-responsive *ONSEN* (*ATCOPIA78*) LTR-RT family in *A. thaliana*. Colors indicate number of TF-binding sites found. GO terms that occur at least six times are highlighted in the plot. All GO-terms and their number of occurrences is listed in **S1 Table**.

**S3 Fig.**
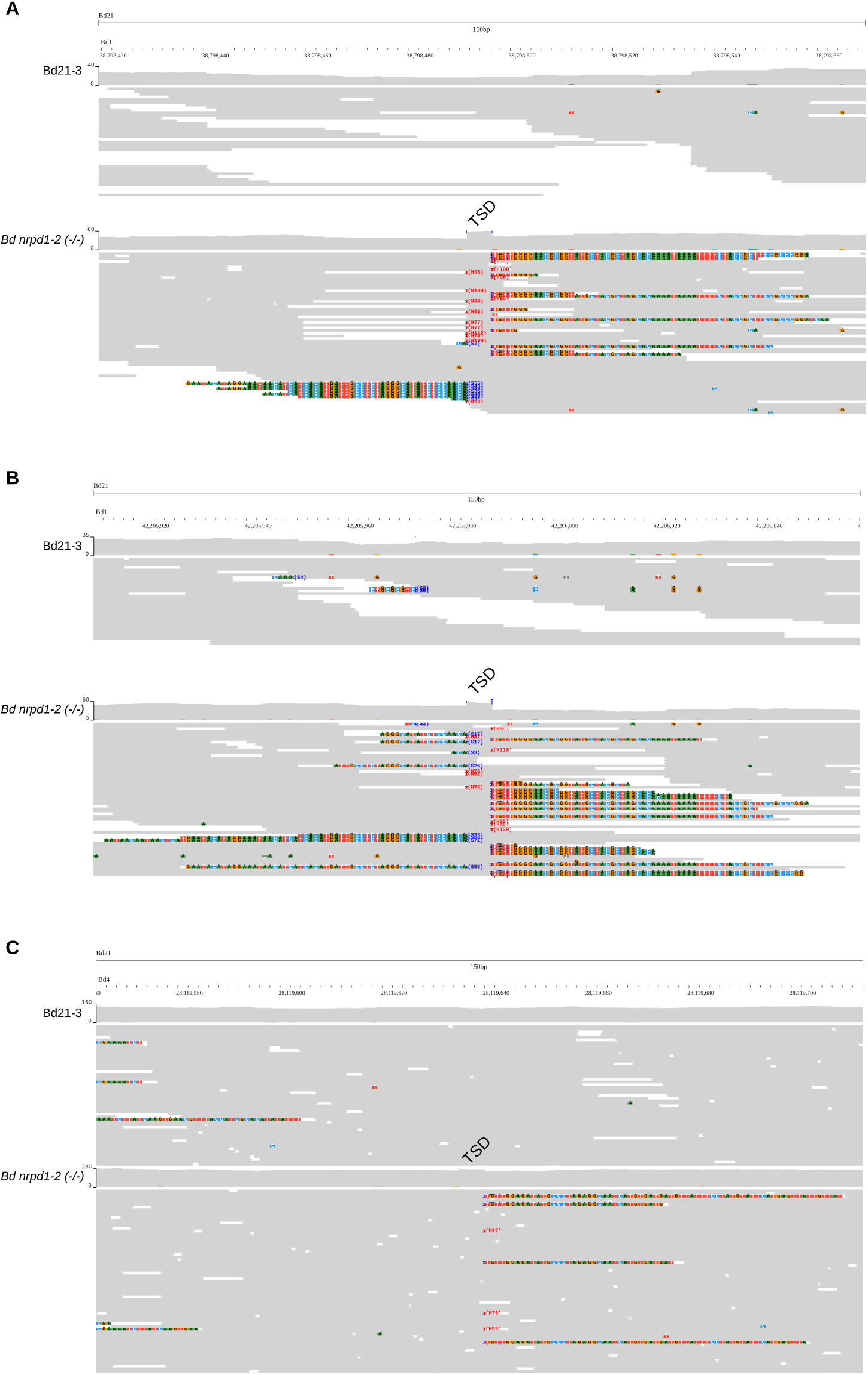
TIPs of *HOPPLA* in one of the resequenced *Bd nrpd1-2 (-/-)* plants. JBrowse screenshot of three insertion sites (**A-C**) in *Bd nrpd1-2 (-/-)* (bottom) compared to the Bd21-3 wt (top). The target side duplication (TSD) is annotated and soft clipped parts of reads are coloured.

**S4 Fig.**
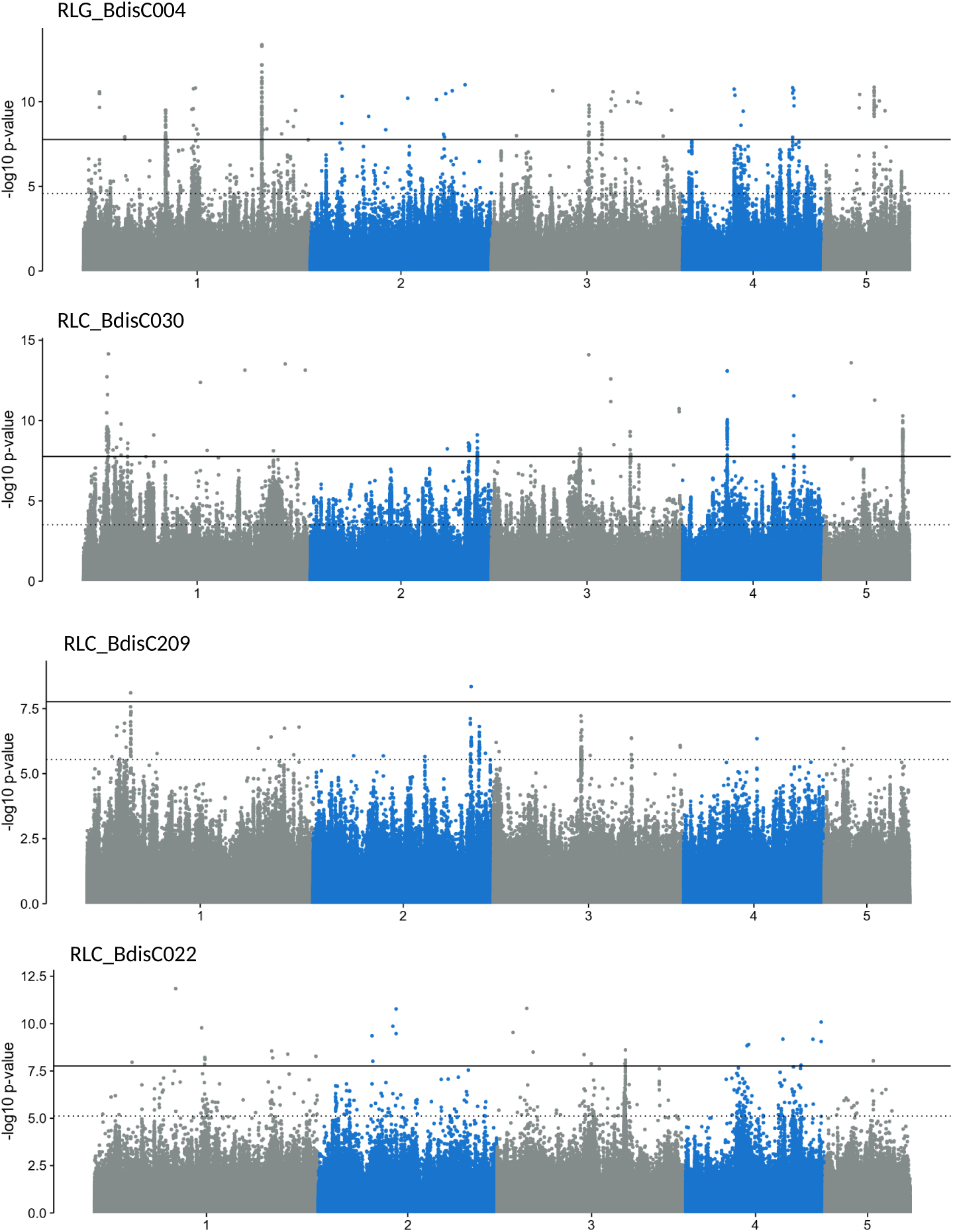
Manhattan plots of the GWASs of pCNs of four recently active LTR-RT families. From top: RLG_BdisC004, RLC_BdisC030 RLC_BdisC209 and RLC_BdisC022. The two significance levels, false discovery rate < 0.05 (dashed line) and Bonferroni correction (solid line) are depicted

## Supporting Information – Table legends

**S1 Table. REVIGO output of the processing of GO terms of transcription factors for which binding sites have been detected in the *HOPPLA* and *ONSEN* consensus sequences.**

**S2 Table. Normalized pCNs of LTR-RTs and bioclimatic variables of the 320 natural accessions of *A. distachyon***

**S3 Table. Gene list of pCNs GWAS with different levels of significance (FDR < 0.05, BC) and window sizes (20 kb, 50 kb)**

**S4 Table. Components of the Pol IV and Pol V holoenzymes in *B. distachyon***

**S5 Table. Sequences of oligos used in this study**

**S6 Table. Meta information of mobilome-seq samples**

